# WEDAP: A Python Package for Streamlined Plotting of Molecular Simulation Data

**DOI:** 10.1101/2024.05.18.594829

**Authors:** Darian T. Yang, Lillian T. Chong

## Abstract

Given the growing interest in path sampling methods for extending the timescales of molecular dynamics (MD) simulations, there has been great interest in software tools that streamline the generation of plots for monitoring the progress of large-scale simulations. Here, we present the WEDAP Python package for simplifying the analysis of data generated from either conventional MD simulations or the weighted ensemble (WE) path sampling method, as implemented in the widely used WESTPA software package. WEDAP facilitates (i) the parsing of WE simulation data stored in highly compressed, hierarchical HDF5 files, and (ii) incorporates trajectory weights from WE simulations into all generated plots. Our Python package consists of multiple user-friendly interfaces: a command-line interface, a graphical user interface, and a Python application programming interface. We demonstrate the plotting features of WEDAP through a series of examples using data from WE and conventional MD simulations that focus on the HIV-1 capsid protein C-terminal domain dimer as a showcase system. The source code for WEDAP is freely available on GitHub at https://github.com/chonglab-pitt/wedap.

## Introduction

To characterize biological processes beyond the timescales of conventional molecular dynamics (cMD) simulations, various enhanced sampling methods have been developed,^1^ including the weighted ensemble (WE) path sampling strategy,^2,3^ which can be carried out using the Weighted Ensemble Simulation Toolkit with Parallelization and Analysis (WESTPA) software package. ^4,5^ WESTPA is a highly scalable, interoperable software package for carrying out WE simulations, with successful applications to the simulation of protein-ligand unbinding,^6^ protein-protein^7^ or protein-DNA binding,^8^ large-scale protein conformational rearrangements,^9^ phase separation of lipid bilayers,^10^ membrane permeability of drug-like molecules,^11^ and protein folding. ^12,13^

To run a WE simulation, WESTPA initiates multiple weighted trajectories (*N*) in parallel from one or more initial conformations (`bstates`). The configurational space is typically divided into bins along a progress coordinate (`pcoord`) towards a target state (`tstate`). After a fixed time interval (*τ*), a resampling procedure is applied in which the trajectory ensemble is evaluated for either splitting (replicating trajectories) or merging (combining trajectories), with the goal of obtaining even coverage of the binned configuration space. This resampling procedure manages and ensures a statistically rigorous conservation of trajectory weights (*w*),where 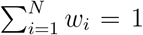. Each WE iteration consists of dynamics propagation for a fixed time interval *τ*, followed by resampling. Trajectories that reach the target state are “recycled” back to the initial state (keeping the same weight) to maintain a non-equilibrium steady state. If trajectories are not recycled, we would refer to the simulation as an equilibrium WE simulation. In addition to the progress coordinate, a number of auxiliary datasets (`auxdata`) can be calculated during a WE simulation for post-simulation analysis. Ideally, the result is an ensemble of unbiased pathways that can be used to directly calculate rate constants between any pair of states.

In this application note, we present the Weighted Ensemble Data Analysis and Plotting (WEDAP; pronounced we-dap) software package for creating plots of varying complexity either from WE or cMD simulation data. WEDAP is currently divided into three submodules: (i) `wedap` for data distributions of WE data, (ii) `mdap` for data distributions of cMD data, and (iii) `wekap` for plotting kinetics data from WE simulations. Each module is available through the command line interface (CLI), a graphical user interface (GUI), or directly through the Python application programming interface (API) (Figure S1). Built upon Matplotlib,^14^ plotting with WEDAP can be useful for tracking simulation progress on remote computing resources and for generating publication-quality figures post-simulation.

WEDAP satisfies the need for an open-source, Python-based software package focused on plotting large-scale simulation data stored in highly compressed, hierarchical HDF5 files (Figure S2). While other software packages for plotting have been developed, these other tools are either not written in Python,^15–17^ limiting API access for custom routines and data pipelines, or are tied to pre-existing analysis packages and specific dynamics engines.^18,19^ For path sampling simulations, there are even fewer plotting tools^20^ available.

### Simulation details

To demonstrate the capabilities of WEDAP, we focus on cMD and WE simulation data from sampling the conformational ensemble of the HIV-1 capsid protein (CA) C-terminal domain (CTD) dimer, herein referred to as CA-CTD (Figure 1). The CTD of the two-domain capsid protein forms a dimer that connects individual chains in the mature capsid lattice. The flexible CA-CTD dimer interface^23^ is a potential target for the development of antiretroviral therapeutics.^24^

**Figure 1:**
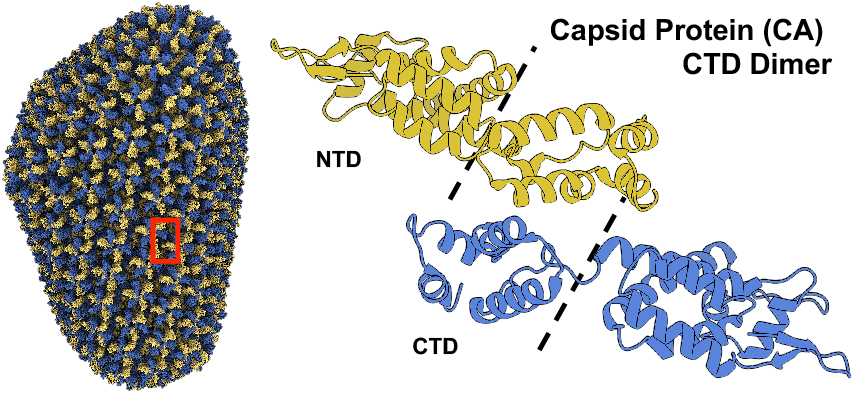
Our plotting examples use either cMD or WE simulation data involving the HIV-1 capsid protein CTD dimer. The CA-CTD dimer connects the subunits of the assembled capsid (PDB ID: 3J3Q^21^), as boxed in red. The full-length capsid protein dimer is expanded and shown using ribbon diagrams (PDB ID: 2M8L^22^), where each monomer is sectioned into the respective N-terminal and C-terminal domains (NTD and CTD) using dashed lines.

Our simulations employed the implicitly polarized AMBER ff15ipq protein force field^25^ with a truncated octahedral box of explicit SPC/E_b_ ^26^ water molecules with a 12 Å clearance between the solute and the edge of the box. Unpaired charges were neutralized by adding Na^+^ or Cl^-^ ions, treated with Joung and Cheatham ion parameters.^27^ Protonation states for ionizable residues were adjusted to represent the major species present at pH 6.5. A 2 fs timestep was enabled in all simulations by constraining all bonds to hydrogen to their equilibrium values using the SHAKE algorithm.^28^

WE simulations were run with a resampling time interval (*τ*) of 100 ps and a 1D progress coordinate tracking the heavy-atom root-mean-square deviation (RMSD). Fixed bins for WE were placed along the progress coordinate at a 0.5 Å interval between 0 Å and 8 Å with a target count of four trajectories per bin. For rate-constant calculations, we used a target state of >5 Å heavy-atom RMSD. Trajectory coordinates were saved every 10 ps for analysis. Overall, we ran 200 WE iterations yielding 0.8 µs of aggregate simulation time. To generate cMD data, we ran a single-µs simulation. The reference structure used for all native-contact and RMSD calculations was the CA-CTD NMR structure (PDB^29^ ID: 2KOD^30^).

### Overview of Examples

Here we present a set of 13 examples demonstrating various features of the WEDAP package. All of our examples use either WE or cMD data from simulations of CA-CTD and the first dimension of the progress coordinate, auxiliary dataset, or cMD dataset. With multi-dimensional inputs, dataset dimensions can be specified using the `--Xindex`, `--Yindex`, or `--Zindex` flags. A Jupyter notebook, along with all corresponding files needed to reproduce each of our examples is available in the WEDAP GitHub repository.

### Example 1: Monitoring the time-evolution of a 1D probability distribution

Our first example generates a plot of the time-evolution of a WE simulation (Figure 2.A), with the probability distribution of the heavy-atom RMSD of CA-CTD on the x-axis and each iteration of the WE simulation on the y-axis. The color bar represents the probabilities of each bin in the histogram. The probability values are derived from the raw counts of all trajectory segments in each WE iteration and appropriately weighted. This weighted histogram is normalized and shown on an inverted natural log scale 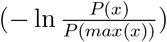, in units of *k*_*B*_*T*. These evolution plots are useful for observing probability changes through the progression of the WE simulation, but are limited to tracking a single dataset. In this example, we use the first dimension of the progress coordinate, but other datasets or dataset dimensions can be used by adjusting the `--Xname` and `--Xindex` flags.

**Figure 2:**
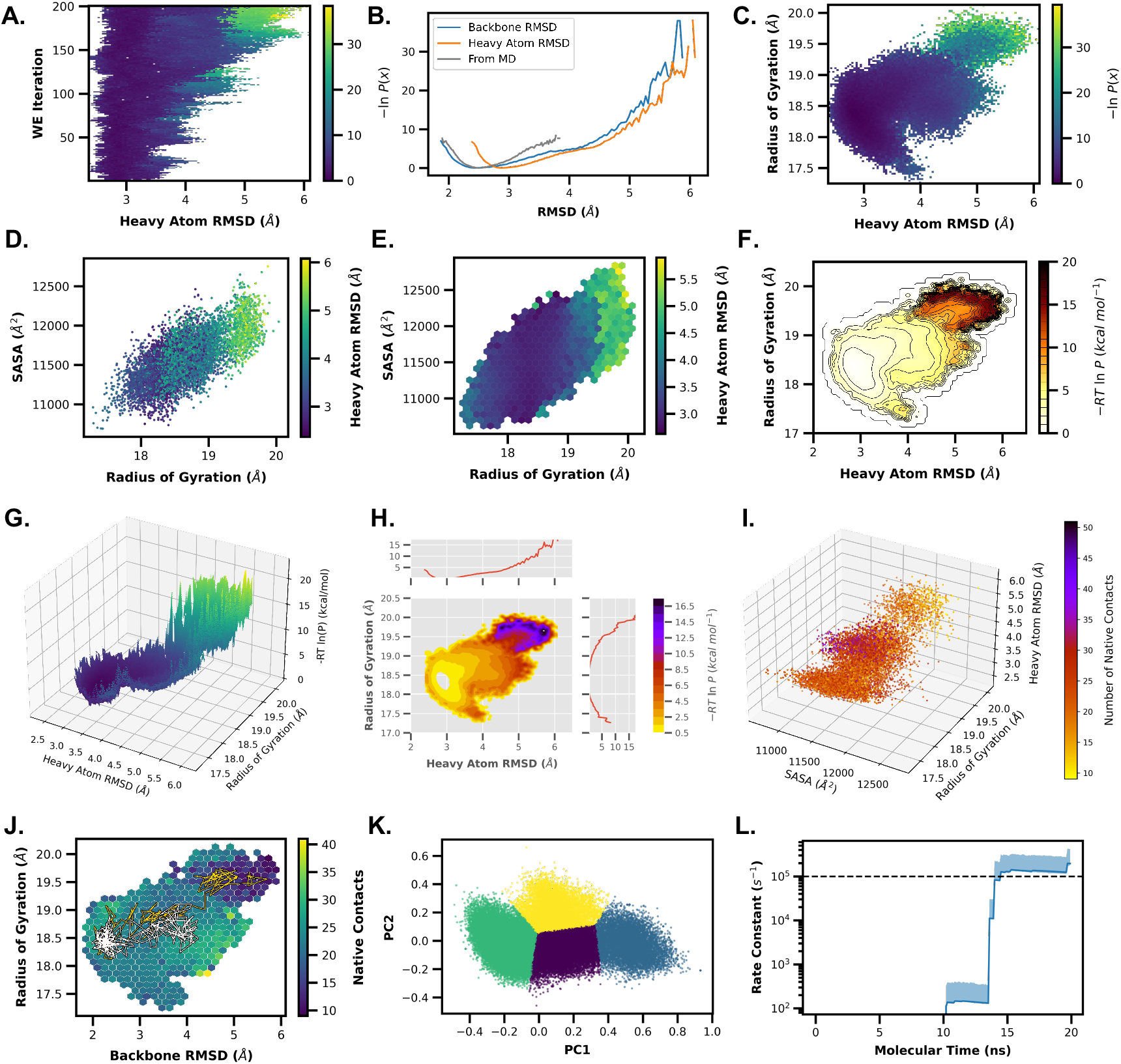
Gallery of WEDAP plots. Each panel of this figure is discussed in the corresponding examples subsection. In brief, we demonstrate how to create plots using WEDAP for 1D WE time-evolutions (A), 1D probability distributions (B), 2D probability distributions (C), 3D scatter plots (D), 3D hexagonal binned plots (E), 2D histograms with contour lines (F), 3D projected contour plots (G), joint plots with contour fills (H), 4D scatter plots (I), trajectory tracing (J), principal component analysis with cluster labels (K), and rate constants over molecular time (L).

### Example 2: Generating a 1D probability distribution of simulation data

We can plot the 1D probability distribution of a single WE iteration, or we can summarize a range of WE iterations. Here, we plot the cumulative probability distribution of the heavy-atom RMSD for the entire range of WE iterations. This allows us to focus on a subset or overview of the data presented in the time-evolution plot (Example 1). The distributions being plotted are summations of histogram counts, normalized and weighted across multiple WE iterations.

We can plot multiple RMSD datasets from a WESTPA HDF5 file using `wedap`, or from cMD simulation data using `mdap` (Figure 2.B). The WE data is represented by a weighted probability distribution while the cMD distribution assumes an equal set of weights.

### Example 3: Generating a 2D probability distribution of simulation data

We can make 2D probability distributions using more than one feature, allowing us to compare both the heavy-atom RMSD and the radius-of-gyration (Figure 2.C) for WE simulation data. These same plots are also available using cMD simulation data.

### Example 4: Generating a scatter plot colored by a feature of interest

Next, we create and color a scatter plot by a feature of interest, such as heavy-atom RMSD. The x-axis data here is the radius-of-gyration and the y-axis data is the solvent-accessible surface area (SASA) (Figure 2.D). Each data point corresponds to a frame (conformation) from the WE or cMD simulation data. The marker size (`--scatter-size`) and amount of data used (`--scatter-interval`) can be customized for clearer visualization.

### Example 5: Generating a hexagonal binned plot colored by a feature of interest

Using the same data from Example 4, we can generate a hexagonal binned plot where each bin is colored by a feature of interest (Figure 2.E). The amount of hexagonal bins can be set using the `--hexbin-grid` flag, and each bin represents the average of all the frames from a WE or cMD simulation that fall within the hexagonal bin boundaries. For WE data, each hexagonal bin is reduced using a weighted average (by default), while for cMD data, a standard, unweighted average is used. By representing our data with hexagonal bins, we avoid issues with overlapping data points that may arise with scatter plots of large datasets.

### Example 6: Generating 2D histograms and contour plots

WEDAP can also create contour plots or combinations of both histograms and contour plots. We again compare heavy-atom RMSD and radius-of-gyration data to generate an initial 2D probability distribution. We set the probability units to 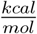 (`--p-units kcal`) with a maximum limit (`--pmax`), and contour lines are overlaid at every single 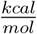 (Figure 2.F).

The histogram uses an alternate color map (`--cmap`), the contour lines are smoothed using a Gaussian noise filter (`--smoothing-level`), contour line-widths are thinned (`--linewidth`), and custom histogram ranges are specified (`--histrange-x` and `--histrange-y`). Likewise, we could compare 2D distributions between multiple simulation datasets from WE or cMD simulations.

### Example 7: Generating 3D projections of contour plots

Using the `--proj3d` or `-3d` flags, we can visualize a contour plot of the heavy-atom RMSD and radius-of-gyration data on a 3D projection (Figure 2.G). Instead of using a flat color bar for the third dimension, seeing the full spatial resolution of the z-axis can be more visually intuitive for interpreting barrier heights, potential metastable states, and pathways between states. We also set a custom contour interval here using the `--contour-interval` flag.

### Example 8: Generating distributions for each dimension of a 2D probability distribution

Joint plots are a useful way to understand both the relationship between two observables in the middle panel (the joint distribution) and the distribution of each observable on the side panels (the marginal distributions). These marginal distributions can be added to any 2D probability distribution from WE or cMD simulation data using the `--joint-plot` or `-jp` flags. Here, we compare the heavy-atom RMSD against the radius-of-gyration using a contour plot with the contour fills only (no contour lines) (Figure 2.H). We include custom color mapping (`--cmap`), probability units (`--p-units`), WE iteration ranges (`--first-iter` and `--last-iter`), probability limits (`--pmin` and `--pmax`), plot style (`--style`), contour data smoothing (`--smoothing-level`), and axes limits (`--xlim` and `--ylim`).

### Example 9: Generating 4D scatter plots

To compare four different features from WE or cMD simulations, we can project a scatter plot onto three dimensions and include a color bar as a fourth dimension (Figure 2.I). This 4D plot is called using the `--proj4d` or `-4d` flags. In this example, we create a 3D scatter plot as a function of SASA, radius-of-gyration, and heavy-atom RMSD. Each data point is colored by the number of native-contacts, as indicated by the color bar. If the color bar is not needed, a 3D projected scatter plot can still be created using the `--proj3d` or `-3d` flags.

### Example 10: Tracing a pathway along a probability distribution of WE data

To trace a single pathway as a function of the WE progress coordinate, we can request, for example, WE iteration 200 and trajectory segment 20 by including `--trace-seg 200 20` in our `wedap` command. Alternatively, we can plot the pathway based on the closest data point to an input set of x- and y-axis values. Using the closest data point, we can trace the WE iteration and trajectory segment pair from, for example, a heavy-atom RMSD of 5.5 Å and radius-of-gyration of 19.5 Å by including `--trace-val 5.5 19.5` in our input `wedap` command. The “trace by value” feature will also output the relevant WE iteration and trajectory segment, enabling easy tracking and further analysis. For this demonstration, we show the trajectory segment-based trace in white and the value-based trace in gold, overlaid on a hexagonal binned plot (Figure 2.J).

### Example 11: Extracting WE data for analysis using external Python libraries

In this next example, we extract the progress coordinate, auxiliary data, and all corresponding trajectory weights using WEDAP. This is done using the Python API, which can then directly interface with other Python libraries such as scikit-learn. ^31^ The WE extracted feature array was then scaled and reduced to two dimensions using principal component analysis. We clustered this reduced feature set using a weighted k-means algorithm, from which the labels were plotted as the colors of each data point along the first two principal components (Figure 2.K). In this manner, we can conveniently extract and use the data from a WESTPA simulation for scikit-learn, PyTorch, ^32^ or any other Python library. This is a useful approach for interfacing WE simulation data with machine learning or deep learning methods.

WEDAP can also be used to make weighted probability distributions along the newly calculated principal components, optionally saving the input dataset into an updated HDF5 file.

### Example 12: Monitoring time-evolution of rate-constant estimates

A primary criterion for monitoring convergence to a non-equilibrium steady state is the leveling off of the rate constant of interest. Here, we use an arbitrary target state with a high heavy-atom RMSD (Figure 2.L). The x-axis is set to either the number of WE iterations or molecular time; defined as *Nτ*, where *N* is the number of WE iterations and *τ* is the fixed time interval of each iteration. The `west.cfg` file used with the `w_ipa` command from WESTPA to generate the resulting `assign.h5` and `direct.h5` files used in this example are provided in the WEDAP GitHub repository.

The `direct.h5` file from WESTPA provides state-to-state flux evolution data. From the Hill relation,^33,34^ we know that in a system with states A and B, probability flux from A to B at steady state is exactly equal to the inverse of the mean first passage time (MFPT):

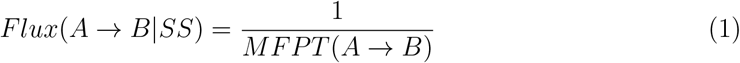

where the MFPT is the average of the first passage times observed during the simulation. If we convert the probability flux units from per *τ* to per second and adjust for concentration dependence (for multi-molecular systems only), we can calculate rate constants over time. For equilibrium WE simulations, the probability flux from A to B is normalized by the state population of A (*Flux*(*A → B*|*SS*)*/P*_*A*_^*eq*^).^35^

Uncertainties for observables estimated using a single simulation are determined as 95% confidence intervals from Monte Carlo block bootstrapping. For multiple simulations, 95% credibility regions are determined using Bayesian bootstrapping.^36^

In this example, we also used a post-processing function (`--postprocess`) to customize our plot from the CLI, adding a horizontal line as an arbitrary reference rate. This customization flag is available with all WEDAP tools and the input is a user-defined Python function.

### Example 13: Creating a time-evolution movie of 2D probability distributions

Because WE data represents an ensemble of trajectories, it is useful to track the time-evolution of more than one dimension or feature. One method to do this is to create a GIF or movie of how two or more features correlate over the course of a WE simulation.

In this example, we use the heavy-atom RMSD and the radius-of-gyration from a CA-CTD WE simulation, looping through the average probability distributions of specified WE iterations (Video S1). We can customize this GIF by setting the range of WE iterations (`--first-iter` and `--last-iter`), optionally setting a larger interval between WE iterations for better performance (`--step-iter`), setting the amount of WE iterations to include in each frame of the GIF (`--avg-plus`), and providing the GIF file output path (`--gif-out`).

### Other WEDAP features

Beyond our set of examples, other notable features of WEDAP include the following: (i) multiple input files for `wedap`, which are automatically normalized and plotted as long as the dataset(s) specified exist in all input H5 files (multiple cMD data files can also be input with `mdap`); (ii) `wedap` and `mdap` input data can be a NumPy^37^ array directly passed to the API, or the name of a file with a `.dat`, `.txt`, `.pkl`, `.npz`, or `.npy` extension; (iii) the WE trajectory ensemble can be filtered in `wedap` to generate probability distributions of only successfully recycled events (`--succ-only`); and (iv) the WE basis states can also be filtered in `wedap` to only plot the probability distributions of trajectories from specific starting states (`--skip-basis`).

With the exception of the WE evolution plot (Figure 2.A), most `wedap` plots can be made using cMD data instead of WE data generated using WESTPA. Although many of the commands and uses for `mdap` were not demonstrated here, examples can be found on the WEDAP demo Jupyter notebook. In future work, we will extend WEDAP to include a module for rate-constant calculations and plots of cMD data, allowing easy comparisons between WE and cMD simulations in terms of the time-evolution of a rate estimate.

## Conclusions

We have presented WEDAP, a software package for plotting observables of interest from both WE and conventional MD simulation data. Based on WE and cMD simulation data from conformational sampling of the HIV-1 capsid protein CTD dimer, we demonstrated the use of WEDAP for generating various plots, including simple 1D probability distributions and animated time evolutions of 2D probability distributions. While we originally implemented WEDAP for generating probability distributions from WE simulations, many additional features have been introduced and ongoing development for other types of plots and simulation datasets is expected to continue.

Overall, WEDAP is designed to provide more accessible data analysis and plotting tools to the simulation community with modularity that allows for flexible usage and easy feature development. Accessible plotting capabilities for many different datasets should facilitate data exploration and simulation monitoring to identify trends and relationships within both WE or cMD data. We note that WEDAP is already being used by the simulation community^13,38,39^ and we expect to continue maintenance and development in parallel with the WESTPA software package. We hope that WEDAP can eventually be adapted to benefit other enhanced sampling communities,^1,40–45^ only a few of which have companion tools for data visualization.^20^

## Notes

LTC serves on the scientific advisory board of OpenEye Scientific Software.

## Data and Software Availability Statement

All plotting tools used are available in the open-source WEDAP software package, with source code available on GitHub: https://github.com/chonglab-pitt/wedap and deposited under DOI: https://doi.org/10.5281/zenodo.11051656. WEDAP is also available on PyPI: https://pypi.org/project/wedap and can be installed using PIP: `pip install wedap`. The open-source WESTPA software package was used to generate the example WE simulation data and is available on GitHub: https://github.com/westpa/westpa. The WESTPA HDF5 data files, cMD data files, and Jupyter notebook needed to reproduce all plotting examples can be found in the WEDAP GitHub repository. Full API documentation and more examples for using WEDAP can be found on the documentation web page: https://darianyang.github.io/wedap.

## Acknowledgement

We would like to thank Jeremy Leung, Riccardo Solazzo, Anthony Bogetti, Marion Silvestrini, Rhea Abraham, and other members of the Chong lab and WESTPA developer community for helpful discussions, testing out WEDAP through various stages, and providing feedback and suggestions.

## Supporting Information Available

We include the following as supporting information:

- Figure S1: Example images of the WEDAP GUI, CLI, and API for `wedap` and `mdap`.
- Figure S2: File organization structure of a typical HDF5 output file from WESTPA.
- Video S1: The animated time-evolution of a 2D probability distribution from Example 13.

## TOC Graphic

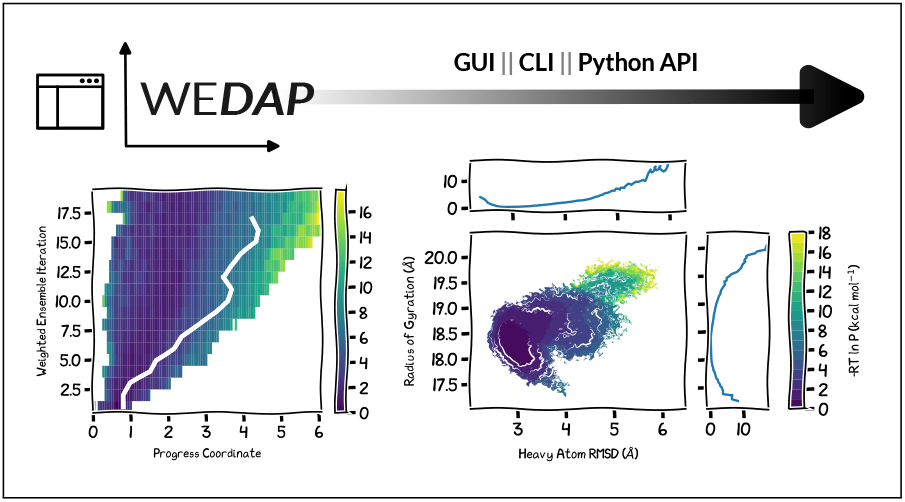

## References

(1) Zwier, M. C.; Chong, L. T. Reaching biological timescales with all-atom molecular dynamics simulations. Current Opinion in Pharmacology 2010, 10, 745–752.

(2) Huber, G. A.; Kim, S. Weighted-ensemble Brownian dynamics simulations for protein association reactions. Biophysical Journal 1996, 70, 97–110.

(3) Zuckerman, D. M.; Chong, L. T. Weighted Ensemble Simulation: Review of Methodology, Applications, and Software. Annual Review of Biophysics 2017, 46, 43–57.

(4) Zwier, M. C.; Adelman, J. L.; Kaus, J. W.; Pratt, A. J.; Wong, K. F.; Rego, N. B.; Suárez, E.; Lettieri, S.; Wang, D. W.; Grabe, M.; Zuckerman, D. M.; Chong, L. T. WESTPA: An interoperable, highly scalable software package for weighted ensemble simulation and analysis. Journal of Chemical Theory and Computation 2015, 11, 800–809.

(5) Russo, J. D. et al. WESTPA 2.0: High-Performance Upgrades for Weighted Ensemble Simulations and Analysis of Longer-Timescale Applications. Journal of Chemical Theory and Computation 2022, acs.jctc.1c01154.

(6) Nunes-Alves, A.; Zuckerman, D. M.; Arantes, G. M. Escape of a Small Molecule from Inside T4 Lysozyme by Multiple Pathways. Biophysical Journal 2018, 114, 1058–1066.

(7) Saglam, A. S.; Chong, L. T. Protein-protein binding pathways and calculations of rate constants using fully-continuous, explicit-solvent simulations. Chemical Science 2019, 10, 2360–2372.

(8) Brossard, E. E.; Corcelli, S. A. Molecular Mechanism of Ligand Binding to the Minor Groove of DNA. J. Phys. Chem. Lett. 2023, 14, 4583–4590.

(9) Sztain, T. et al. A glycan gate controls opening of the SARS-CoV-2 spike protein. Nature Chemistry 2021 13:10 2021, 13, 963–968.

(10) Poruthoor, A. J.; Sharma, A.; Grossfield, A. Understanding the free-energy landscape of phase separation in lipid bilayers using molecular dynamics. Biophysical Journal 2023, 122, 4144–4159.

(11) Zhang, S.; Thompson, J. P.; Xia, J.; Bogetti, A. T.; York, F.; Skillman, A. G.; Chong, L. T.; LeBard, D. N. Mechanistic Insights into Passive Membrane Permeability of Drug-like Molecules from a Weighted Ensemble of Trajectories. J. Chem. Inf. Model. 2022, 62, 1891–1904.

(12) Adhikari, U.; Mostofian, B.; Copperman, J.; Subramanian, S. R.; Petersen, A. A.; Zuckerman, D. M. Computational Estimation of Microsecond to Second Atomistic Folding Times. 141, 6519–6526.

(13) Santhouse, J. R.; Leung, J. M. G.; Chong, L. T.; Horne, W. S. Effects of altered backbone composition on the folding kinetics and mechanism of an ultrafast-folding protein. Chem. Sci. 2024, 15, 675–682.

(14) Hunter, J. D. Matplotlib: A 2D Graphics Environment. Computing in Science & Engineering 2007, 9, 90–95.

(15) Margreitter, C.; Oostenbrink, C. MDplot: Visualise Molecular Dynamics. R J 2017, 9, 164–186.

(16) Turner, P. J.; Team, G. D. XMGRACE: Version 5.1.22. 2008.

(17) Williams, T.; Kelley, C.; many others, Gnuplot 6.0: an interactive plotting program. http://gnuplot.sourceforge.net/, 2023.

(18) Yekeen, A. A.; Durojaye, O. A.; Idris, M. O.; Muritala, H. F.; Arise, R. O. CHAPERONg : A tool for automated GROMACS-based molecular dynamics simulations and trajectory analyses. Computational and Structural Biotechnology Journal 2023, 21, 4849–4858.

(19) Maity, D.; Pal, D. MD DaVis: interactive data visualization of protein molecular dynamics. Bioinformatics 2022, 38, 3299–3301.

(20) Aarøen, O.; Kiær, H.; Riccardi, E. PyVisA: Visualization and Analysis of path sampling trajectories. Journal of Computational Chemistry 2021, 42, 435–446.

(21) Zhao, G.; Perilla, J. R.; Yufenyuy, E. L.; Meng, X.; Chen, B.; Ning, J.; Ahn, J.; Gronenborn, A. M.; Schulten, K.; Aiken, C.; Zhang, P. Mature HIV-1 capsid structure by cryo-electron microscopy and all-atom molecular dynamics. Nature 2013, 497, 643–646.

(22) Deshmukh, L.; Schwieters, C. D.; Grishaev, A.; Ghirlando, R.; Baber, J. L.; Clore, G. M. Structure and Dynamics of Full-Length HIV-1 Capsid Protein in Solution. J. Am. Chem. Soc. 2013, 135, 16133–16147.

(23) Byeon, I.-J. L.; Hou, G.; Han, Y.; Suiter, C. L.; Ahn, J.; Jung, J.; Byeon, C.-H.; Gronenborn, A. M.; Polenova, T. Motions on the Millisecond Time Scale and Multiple Conformations of HIV-1 Capsid Protein: Implications for Structural Polymorphism of CA Assemblies. J. Am. Chem. Soc. 2012, 134, 6455–6466.

(24) McFadden, W. M.; Snyder, A. A.; Kirby, K. A.; Tedbury, P. R.; Raj, M.; Wang, Z.; Sarafianos, S. G. Rotten to the core: antivirals targeting the HIV-1 capsid core. 18, 41.

(25) Debiec, K. T.; Cerutti, D. S.; Baker, L. R.; Gronenborn, A. M.; Case, D. A.; Chong, L. T. Further along the Road Less Traveled: AMBER ff15ipq, an Original Protein Force Field Built on a Self-Consistent Physical Model. J. Chem. Theory Comput. 2016, 12, 3926–3947.

(26) Takemura, K.; Kitao, A. Water Model Tuning for Improved Reproduction of Rotational Diffusion and NMR Spectral Density. J. Phys. Chem. B 2012, 116, 6279–6287.

(27) Joung, I. S.; Cheatham, T. E. I. Determination of Alkali and Halide Monovalent Ion Parameters for Use in Explicitly Solvated Biomolecular Simulations. J. Phys. Chem. B 2008, 112, 9020–9041.

(28) Ryckaert, J.-P.; Ciccotti, G.; Berendsen, H. J. C. Numerical integration of the cartesian equations of motion of a system with constraints: molecular dynamics of n-alkanes. Journal of Computational Physics 1977, 23, 327–341.

(29) Berman, H. M.; Westbrook, J.; Feng, Z.; Gilliland, G.; Bhat, T. N.; Weissig, H.; Shindyalov, I. N.; Bourne, P. E. The Protein Data Bank. Nucleic Acids Research 2000, 28, 235–242.

(30) Byeon, I.-J. L.; Meng, X.; Jung, J.; Zhao, G.; Yang, R.; Ahn, J.; Shi, J.; Concel, J.; Aiken, C.; Zhang, P.; Gronenborn, A. M. Structural Convergence between Cryo-EM and NMR Reveals Intersubunit Interactions Critical for HIV-1 Capsid Function. Cell 2009, 139, 780–790.

(31) Pedregosa, F. et al. Scikit-learn: Machine Learning in Python. Journal of Machine Learning Research 2011, 12, 2825–2830.

(32) Paszke, A. et al. PyTorch: An Imperative Style, High-Performance Deep Learning Library. Advances in Neural Information Processing Systems. 2019.

(33) Hill, T. Free Energy Transduction and Biochemical Cycle Kinetics; Dover Publications, 2004.

(34) Bhatt, D.; Zuckerman, D. M. Beyond Microscopic Reversibility: Are Observable Nonequilibrium Processes Precisely Reversible? J. Chem. Theory Comput. 2011, 7, 2520–2527.

(35) Suárez, E.; Lettieri, S.; Zwier, M. C.; Stringer, C. A.; Subramanian, S. R.; Chong, L. T.; Zuckerman, D. M. Simultaneous Computation of Dynamical and Equilibrium Information Using a Weighted Ensemble of Trajectories. J. Chem. Theory Comput. 2014, 10, 2658–2667.

(36) Mostofian, B.; Zuckerman, D. M. Statistical Uncertainty Analysis for Small-Sample, High Log-Variance Data: Cautions for Bootstrapping and Bayesian Bootstrapping. J. Chem. Theory Comput. 2019, 15, 3499–3509.

(37) Harris, C. R. et al. Array programming with NumPy. Nature 2020, 585, 357–362.

(38) Bogetti, X.; Bogetti, A.; Casto, J.; Rule, G.; Chong, L.; Saxena, S. Direct observation of negative cooperativity in a detoxification enzyme at the atomic level by Electron Paramagnetic Resonance spectroscopy and simulation. 32, e4770.

(39) Bogetti, A. T.; Leung, J. M. G.; Chong, L. T. LPATH: A Semiautomated Python Tool for Clustering Molecular Pathways. 63, 7610–7616.

(40) Allen, R. J.; Valeriani, C.; Wolde, P. R. t. Forward flux sampling for rare event simulations. J. Phys.: Condens. Matter 2009, 21, 463102.

(41) Swenson, D. W. H.; Prinz, J.-H.; Noe, F.; Chodera, J. D.; Bolhuis, P. G. OpenPath-Sampling: A Python Framework for Path Sampling Simulations. 1. Basics. J. Chem. Theory Comput. 2019, 15, 813–836.

(42) Swenson, D. W. H.; Prinz, J.-H.; Noe, F.; Chodera, J. D.; Bolhuis, P. G. OpenPath-Sampling: A Python Framework for Path Sampling Simulations. 2. Building and Customizing Path Ensembles and Sample Schemes. J. Chem. Theory Comput. 2019, 15, 837–856.

(43) Elber, R. Milestoning: An Efficient Approach for Atomically Detailed Simulations of Kinetics in Biophysics. Annual Review of Biophysics 2020, 49, 69–85.

(44) Warmflash, A.; Bhimalapuram, P.; Dinner, A. R. Umbrella sampling for nonequilibrium processes. The Journal of Chemical Physics 2007, 127, 154112.

(45) Vervust, W.; Zhang, D. T.; Ghysels, A.; Roet, S.; van Erp, T. S.; Riccardi, E. PyRETIS 3: Conquering rare and slow events without boundaries. Journal of Computational Chemistry 2024, 45, 1224–1234.

