## Supplemental Information for "WEDAP: A Python Package for Streamlined Plotting of Molecular Simulation Data"

### Supporting Figures

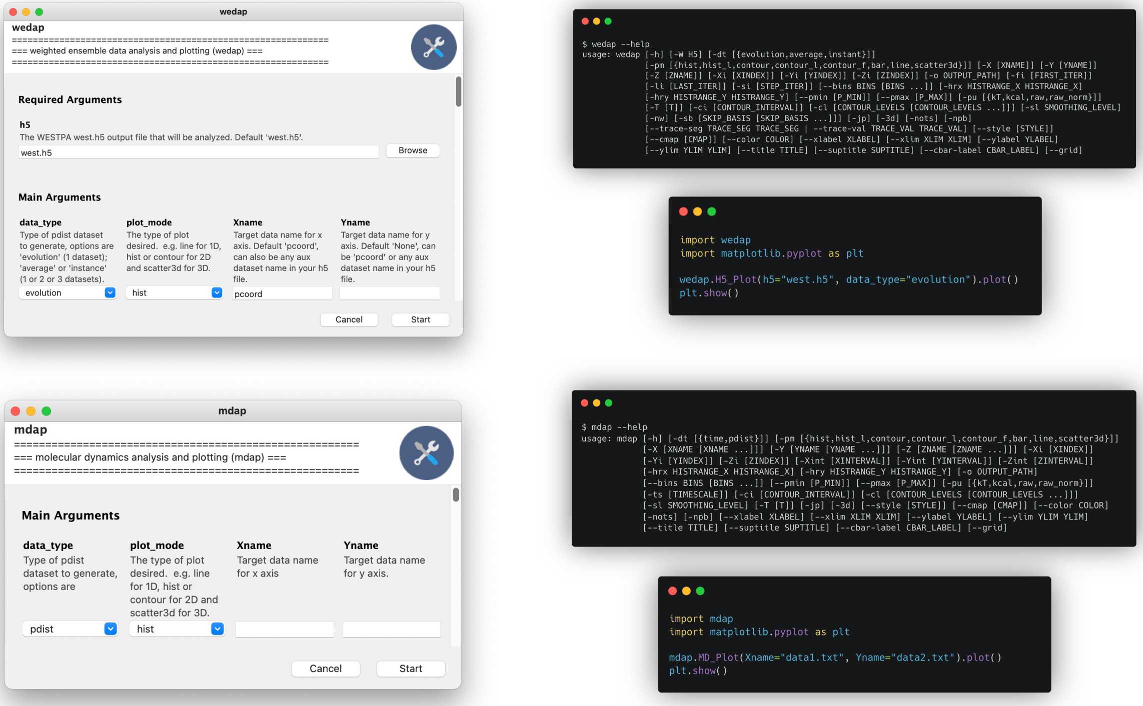


Figure S1: Within the WEDAP toolkit, `wedap`, `mdap`, and `wekap` are all available to use through a graphical user interface, the command line, or using the Python API. An example of each is shown for `wedap` (top) and `mdap` (bottom). The respective interfaces for `wekap` are not shown for simplicity.

```
/
    ibstates/
        index
        naming
            bstate_index
            bstate_pcoord
            istate_index
            istate_pcoord
    tstates/
        index
    bin_topologies/
        index
        pickles
    iterations/
        iter_XXXXXXXX/
            auxdata/
            bin_target_counts
            ibstates/
                bstate_index
                bstate_pcoord
                istate_index
                istate_pcoord
            trajectories
            pcoord
            seg_index
            wtgraph
        ...
    summary
```

Figure S2: The file organization structure of the output H5 file from WESTPA.
